# Hepatitis C virus NS3/4A protease cleaves SPG20, a key regulator of lipid droplet turnover, to promote lipid droplet formation

**DOI:** 10.1101/2025.05.21.655255

**Authors:** Chieko Matsui, Putu Yuliandari, Lin Deng, Takayuki Abe, Ikuo Shoji

## Abstract

Hepatitis C virus (HCV) assembles in close proximity to lipid droplets (LDs), which play important roles in HCV RNA replication. HCV infection often causes the accumulation of large LDs in hepatocytes. However, the molecular mechanism underlying HCV-induced large LD formation is poorly understood. It has been reported that the SPG20/Spartin protein associates with the LD surface and plays a crucial role in LD turnover by recruiting the ubiquitin ligase Itch to promote the ubiquitin-dependent degradation of adipophilin (ADRP), which protects LDs from lipase-mediated degradation. To elucidate the mechanism underlying HCV-induced large LD formation, we investigated the SPG20 protein’s role in LD formation in HCV J6/JFH1-infected Huh-7.5 cells. Immunoblot analysis revealed that HCV infection promoted SPG20 protein cleavage. Transfection of increasing amounts of NS3/4A, but not the inactive NS3/4A mutant, resulted in SPG20 cleavage, implicating the NS3/4A protease in this cleavage. Site-directed mutagenesis suggested that the NS3/4A protease cleaves SPG20 at Cys^504^ and Cys^562^. The SPG20 protein was co-immunoprecipitated with the LD-attached protein TIP47. Increasing amounts of NS3/4A protease, but not inactive NS3/4A, decreased the co-precipitation of SPG20 with TIP47. The siRNA-mediated knockdown of Itch in Huh-7.5 cells restored ADRP levels, suggesting that Itch mediates ubiquitylation-dependent ADRP degradation. Immunofluorescence staining of HCV-infected cells revealed that ADRP was localized mainly around LDs in HCV-infected cells, whereas cytosolic ADRP was decreased. We propose that the HCV NS3/4A protease specifically cleaves SPG20 and inhibits Itch-mediated ubiquitin-dependent degradation of LD-associated ADRP, thereby promoting the formation of large LDs.

**IMPORTANCE:** HCV infection often promotes the formation of large LDs in HCV-infected cells. However, the molecular mechanism underlying large LD formation is poorly understood. LD turnover is regulated by SPG20, Itch E3 ligase, and ADRP. To elucidate the mechanism underlying the formation of large LDs induced by HCV infection, we investigated the roles of SPG20, Itch, and ADRP in large LD formation. The HCV NS3/4A protease specifically cleaves SPG20 and disrupts Itch recruitment to LD-associated ADRP. Therefore, LD-associated ADRP can escape and protects LDs from lipase-mediated degradation, thereby promoting LD growth. We propose that HCV NS3/4A protease-mediated cleavage of SPG20 contributes to a previously uncharacterized mechanism underlying the formation of large LDs in HCV-infected cells. These findings may lead to a better understanding of how the virus forms large LDs in infected cells.

## INTRODUCTION

Hepatitis C virus (HCV) is a positive-sense, single-stranded RNA virus belonging to the *Hepacivirus* genus of the *Flaviviridae* family (1). HCV infection often causes chronic hepatitis, liver cirrhosis and hepatocellular carcinoma (HCC). The HCV genome consists of a 9.6kb RNA encoding a polyprotein of 3,010 amino acids (aa). The polyprotein is cleaved co-translationally and post-translationally into at least 10 proteins by viral proteases and cellular signalases. The viral proteins required for RNA replication include NS3, NS4A, NS4B, NS5A and NS5B. The NS3/4A serine protease is responsible for the processing of HCV proteins (1). Highly potent direct-acting antivirals (DAAs) against the NS3/4A protease, NS5A protein and NS5B polymerase have been developed, representing a breakthrough in HCV therapy (2). DAA therapy results in a >90% sustained virological response (SVR) rate. However, the occurrence of HCC following SVR in patients has been reported, and the molecular mechanism underlying post-SVR HCC is poorly understood (3).

Fatty liver is observed more often in hepatitis C patients than in the general population or in hepatitis B patients (4). These clinical findings suggest that HCV infection may directly cause fatty changes in patients’ hepatocytes. However, the molecular mechanisms underlying HCV-induced fatty changes are poorly understood (5, 6). Lipid droplets (LDs) are intracellular organelles that consists of cholesterol esters and triacylglycerol. LDs in hepatocytes play important roles in HCV RNA replication and the production of infectious HCV (7–9). LDs are associated with phospholipid membranes and the PAT family proteins. The PAT family of protein consists of perilipin (PLIN1), adipophilin (ADRP/PLIN2), tail-interacting protein of 47 kDa (TIP47/PLIN3), S3-12 (PLIN4), and MLDP (PLIN5/ OXPAT) (10, 11). These proteins are important for the biogenesis, stabilization and degradation of LDs (12). Among them, ADRP is the most abundant LD-associated protein in hepatocytes (13).

SPG20, which is also known as Spartin, plays an important role in LD turnover (14–16). SPG20 was initially identified as the target gene of autosomal recessive hereditary spastic paraplegia in Troyer syndrome (17). Frameshift mutations in the SPG20 gene cause Troyer syndrome. SPG20 harbors three domains: a microtubule-interacting and trafficking domain (MIT), a plant-senescence domain (PSD), and a ubiquitin-binding region (UBR) (18). The SPG20 PSD domain is responsible for its association with LDs (14).

SPG20 interacts with several Nedd4 family ubiquitin ligases, such as atrophin-1-interacting protein 4 (AIP4/Itch) (15, 19), atrophin-1-interacting protein 5, (AIP5/WWP1) (14, 15) and atrophin-1-interacting protein 2, (AIP2/WWP2). Nedd4 family ubiquitin ligases possess an N-terminal C2 domain responsible for subcellular localization and two to four WW domains that are important for binding to the substrates (20). The association of SPG20 with an adaptor protein promotes ubiquitin ligase activity (21). SPG20 interacts with Itch via protein-protein interactions between the WW domain of Itch and the PPxY motif of SPG20. SPG20 recruits Itch to ADRP, resulting in the ubiquitylation and degradation of ADRP. ADRP protects LDs from lipase-mediated degradation, whereas degradation of ADRP results in disruption of growth of LDs (22). We previously reported that HCV infection induces JNK activation (23, 24). We also reported that HCV-induced JNK activation results in Itch phosphorylation and activation (25).

In this study, we sought to identify the previously uncharacterized mechanism underlying the formation of large LDs in HCV-infected cells. We provided evidence that the HCV NS3/4A protease specifically cleaves SPG20 and inhibits the Itch-mediated ubiquitin-dependent degradation of LD-associated ADRP, thereby promoting the formation of large LDs.

## RESULTS

### HCV infection increases the size and diameter but not the number of LDs

To examine the size and number of LDs in HCV-infected cells, we performed immunofluorescence staining using a BODIPY lipid probe. Immunofluorescence staining revealed that the LDs in HCV J6/JFH1-infected Huh-7.5 cells (Fig. 1A, 2nd panel, left) were larger than those in mock-infected cells (Fig. 1A, first panel, left). To determine the role of HCV infection in LD formation, we treated the cells with the NS3/4A protease inhibitor VX950. Treatment of the cells with VX950 resulted in a reduction in LD size (Fig. 1A, third panel, left). These results suggest that HCV infection increases LD size.

**Fig. 1.**
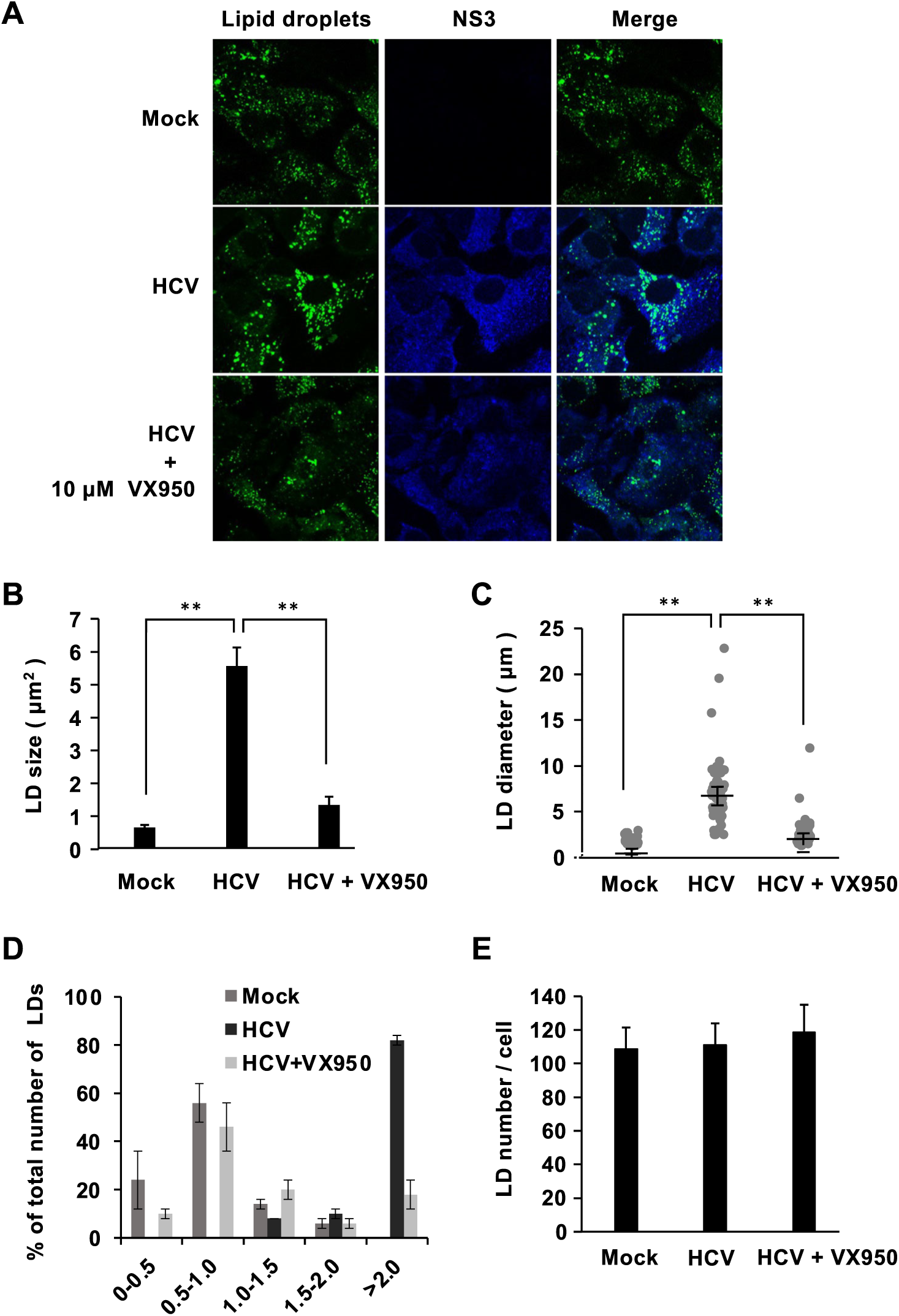
HCV infection increased the size of LDs, but not the number of LDs. (A-E) Huh-7.5 cells were plated at 1.0×10^5^ cells per 24-well plate and cultured for 12 h. The cells were infected with HCV J6/JFH1 at a multiplicity of infection (m.o.i.) of 2. The cells were cultured in the presence or absence of the NS3/4A inhibitor for 48 h as indicated. At 48 h post-infection, the cells were fixed with 4% paraformaldehyde and stained with anti-NS3 mouse MAb followed by Alexa Fluor 405-conjugated goat anti-mouse IgG (blue). LDs were stained with BODIPY493/503 (green). The stained cells were examined under a confocal laser scanning microscope. The diameters of the individual BODIPY-positive areas were measured via the ImageJ software. The results are given from three independent experiments. Fifty cells were counted in each condition. (B) Measurements of LD sizes. (C) Measurements of LD diameter distribution. Individual BODIPY-positive areas were visualized. (D) Measurements of LD size distribution. (E) Numbers of LDs in Huh-7.5 cells. Individual BODIPY-positive areas were counted to determine the number of LDs. **, P<0.01, compared with HCV-infected cells.

We used ImageJ software to measure LD size and diameter. We observed that the LDs in HCV-infected cells were larger and had greater diameters than those in mock-infected cells (Figs. 1B, 1C, and 1D). On the other hand, treatment of the cells with the NS3/4A inhibitor VX950 reduced LD size and diameter. We counted the LDs in HCV-infected cells and in mock-infected cells via a BODIPY lipid probe under a microscope. The number of LDs per cells was not increased in HCV-infected cells (Fig. 1E). These results suggest that HCV infection increases the size and diameter of LDs, but not their number.

### HCV NS3/4A protease activity is responsible for the cleavage of SPG20

To determine the effects of HCV-infection on the SPG20 protein, which is known as a key regulator of LD turnover, we performed immunoblot analysis of the SPG20 protein in HCV-infected Huh-7.5 cells. The SPG20 band decreased and the short form of SPG20 was detected in HCV-infected cells at 3, 5, and 7-days post-infection (Fig. 2A, upper panel, lanes, 6, 8, and 10). This result suggested that SPG20 was cleaved in HCV-infected cells.

**Fig. 2.**
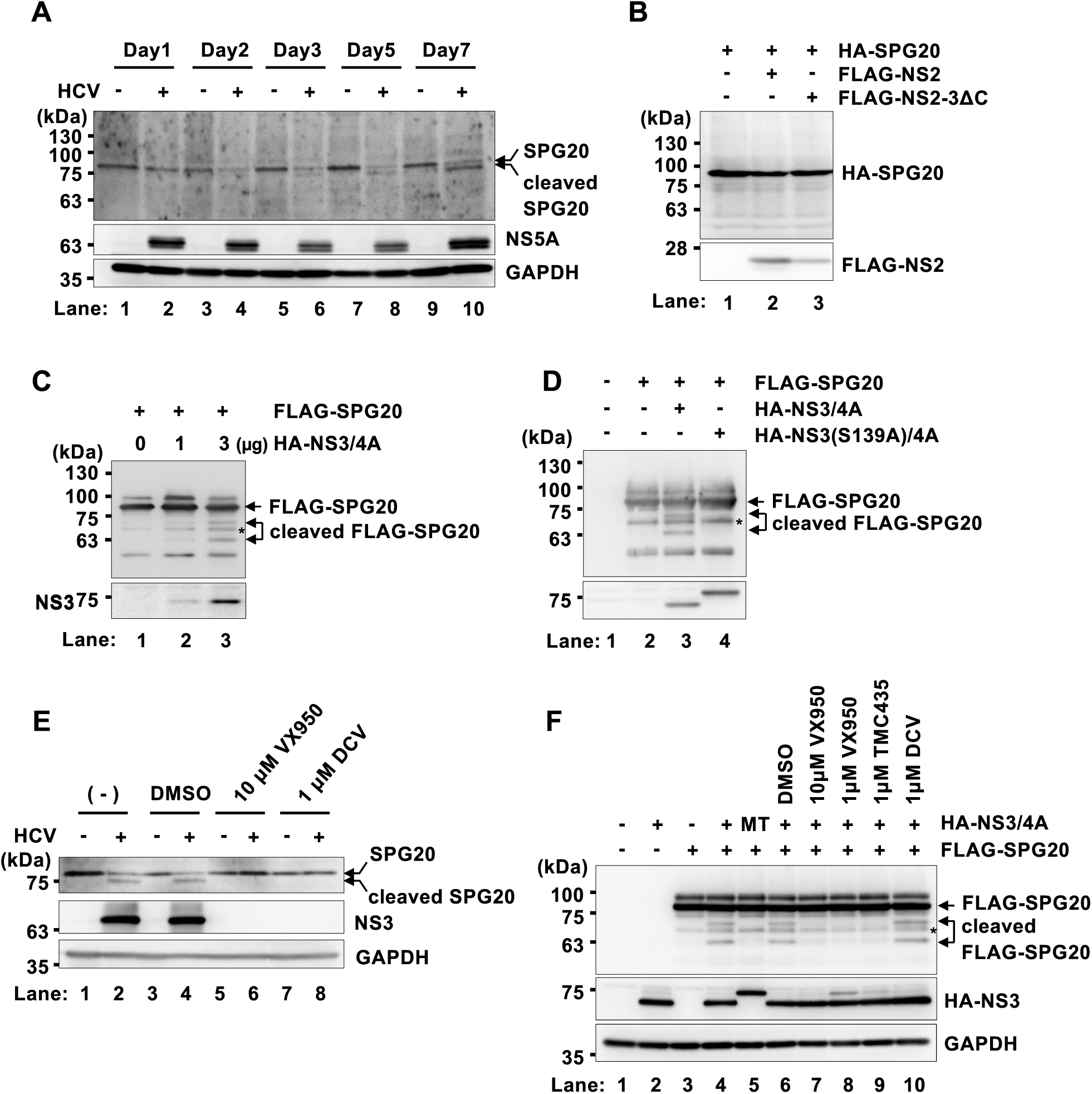
HCV NS3/4A protease activity is responsible for the cleavage of SPG20. (A) Huh-7.5 cells were plated in 2.0×10^5^ cells per 6-well plate and cultured for 12 h. The cells were infected with HCV J6/JFH1 at an m.o.i. of 2. The cells were cultured and harvested at the indicated time points. The cells were analyzed by immunoblotting with anti-SPG20 goat PAb, anti-NS5A rabbit PAb and anti-GAPDH mouse MAb. The level of GAPDH served as a loading control. (B) Huh-7.5 cells were plated at 1.7×10^6^ cells per 10 cm dish and cultured for 12 h. The cells were subsequently transfected with pCAG-HA-SPG20 together with either pCAG-FLAG-NS2 or pCAG-FLAG-NS2-3ΔC. At 48 h after transfection, the cells were harvested. Whole cell lysates were analyzed by immunoblotting with anti-HA rabbit PAb and anti-FLAG M2 mouse MAb. (C) Huh-7.5 cells were plated at 1.7×10^6^ cells per 10 cm dish and cultured for 12 h. The cells were then transfected with pCAG-FLAG-SPG20 together with pCAG-HA-NS3/4A. At 48 h after transfection, the cells were harvested. The cell lysates were immunoprecipitated with anti-FLAG M2 beads, and the bound proteins were immunoblotted with anti-FLAG MAb and anti-HA PAb. *, non-specific band. (D) Huh-7.5 cells were plated at 1.7×10^6^ cells per 10 cm dish and cultured for 12 h. The cells were subsequently transfected with pCAG-FLAG-SPG20 together with either pCAG-HA-NS3/4A or pCAG-HA-NS3(S139A)/4A. At 48 h after transfection, the cells were harvested. The cell lysates were immunoprecipitated with anti-FLAG M2 beads, and the bound proteins were immunoblotted with anti-FLAG MAb and anti-HA PAb. *, non-specific band. (E) Huh-7.5 cells were plated in 2.0×10^5^ per 6-well plates and cultured for 12 h. The cells were infected with HCV J6/JFH1 at an m.o.i. of 2, and cultured for 6 days. The cells were treated with an NS3 inhibitor (VX950) or an NS5A inhibitor (daclatasvir) for 6 days. At 6 days post-infection, the cells were harvested and analyzed by immunoblotting with anti-SPG20 PAb, anti-NS3 MAb and anti-GAPDH MAb. The level of GAPDH served as a loading control. (F) Huh-7.5 cells were transfected with pCAG-FLAG-SPG20 together with either pCAG-HA-NS3/4A or pCAG-HA-NS3(S139A)/4A. The cells were treated with the NS3 inhibitor (VX950 or TMC435) or the NS5A inhibitor (daclatasvir) for 48 h. At 48 h posttransfection, the cells were harvested and analyzed by immunoblotting with anti-FLAG MAb, anti-HA PAb and anti-GAPDH MAb. The level of GAPDH served as a loading control. *, non-specific band.

To examine whether HCV proteases are involved in the cleavage of SPG20, we examined the effects of either the HCV NS2 protease or the NS3/4A protease on the cleavage of SPG20. We transfected cells with pCAG-HA-SPG20 together with either pCAG-FLAG-NS2 or pCAG-FLAG-NS2-3ΔC. Immunoblot analysis revealed that FLAG-NS2 was processed in the cells transfected with pCAG-FLAG-NS2-3ΔC, suggesting that the NS2 protease was active and cleaved NS2-3ΔC at the NS2/NS3 cleavage site (Fig. 2B, lower panel, lane 3). However, the short form of SPG20 was not detected in the presence of the HCV NS2 protease (Fig. 2B, upper panel, lane 3), suggesting that the HCV NS2 protease is not responsible for the cleavage of SPG20. To determine whether the NS3/4A protease is involved in the cleavage of SPG20, Huh-7.5 cells were transfected with FLAG-SPG20 and increasing amounts of HA-NS3/4A. The overexpression of HA-NS3/4A resulted in the appearance of short forms of SPG20, suggesting that the HCV NS3/4A protease plays a role in the cleavage of SPG20 (Fig. 2C, upper panel, lane 3). To determine whether SPG20 is cleaved by the NS3/4A protease, FLAG-SPG20 was co-expressed with either HA-NS3/4A or the inactive mutant HA-NS3(S139A)/4A in Huh-7.5 cells. Immunoblot analysis demonstrated that cleavage of SPG20 occurred only in the presence of wild-type NS3/4A and not in the presence of HA-NS3(S139A)/4A (Fig. 2D, upper panel, lanes 3 and 4). These results suggest that HCV NS3/4A protease activity is responsible for the cleavage of SPG20.

### HCV-induced cleavage of SPG20 is inhibited by the NS3/4A protease inhibitor

To determine whether the NS3/4A protease inhibitor inhibits the HCV-induced cleavage of SPG20, we examined the effects of the specific NS3/4A protease inhibitor VX950 on the cleavage of SPG20. Huh-7.5 cells were infected with HCV J6/JFH1 with or without VX950 treatment. When the cells were treated with VX950, the cleaved form of SPG20 disappeared (Fig. 2E, upper panel, lane 6). We also treated the cells with the NS5A inhibitor daclatasvir (DCV). Treatment of HCV-infected cells with DCV reduced the level of cleaved SPG20, presumably due to the suppression of HCV replication (Fig. 2E, upper panel, lane 8). We then examined the cleavage of exogenous SPG20 by NS3/4A in a transient expression system. Huh-7.5 cells were co-transfected with the HA-NS3/4A plasmid and the FLAG-SPG20 plasmid, and the cells were treated with the NS3/4A protease inhibitor VX950 or TMC435. The cleavage of exogenous SPG20 was inhibited by either VX950 or TMC435 (Fig. 2F, upper panel, lanes 7, 8, and 9), whereas DCV had no inhibitory effect on the cleavage of SPG20 (Fig. 2F, upper panel, lane 10). These results indicate that HCV-induced cleavage of SPG20 is dependent on NS3/4A protease activity.

### NS3/4A protease cleaves SPG20 at Cys^504^ and Cys^562^

To determine the cleavage sites on SPG20 by the NS3/4A protease, we introduced substitution mutations at possible cleavage sites on SPG20. There were at least four possible cleavage sites on SPG20 by HCV NS3/4A, considering the consensus sequence of the NS3/4A protease cleavage site (Fig. 3B, bold, underlined). To detect cleaved SPG20, we constructed a plasmid expressing SPG20 as a C-terminally FLAG-tagged protein. The cells expressed either SPG20-FLAG or HA-NS3/4A, and the cell lysates were immunoprecipitated with FLAG M2 beads. Immunoprecipitation analysis revealed two cleaved SPG20 bands with molecular masses of approximately 22 kDa and 19 kDa (Fig. 3C, upper panel, lane 4, arrows) in the presence of HCV NS3/4A but not in the presence of HCV NS3(S139A)/4A (Fig. 3C, upper panel, lane 5).

**Fig. 3.**
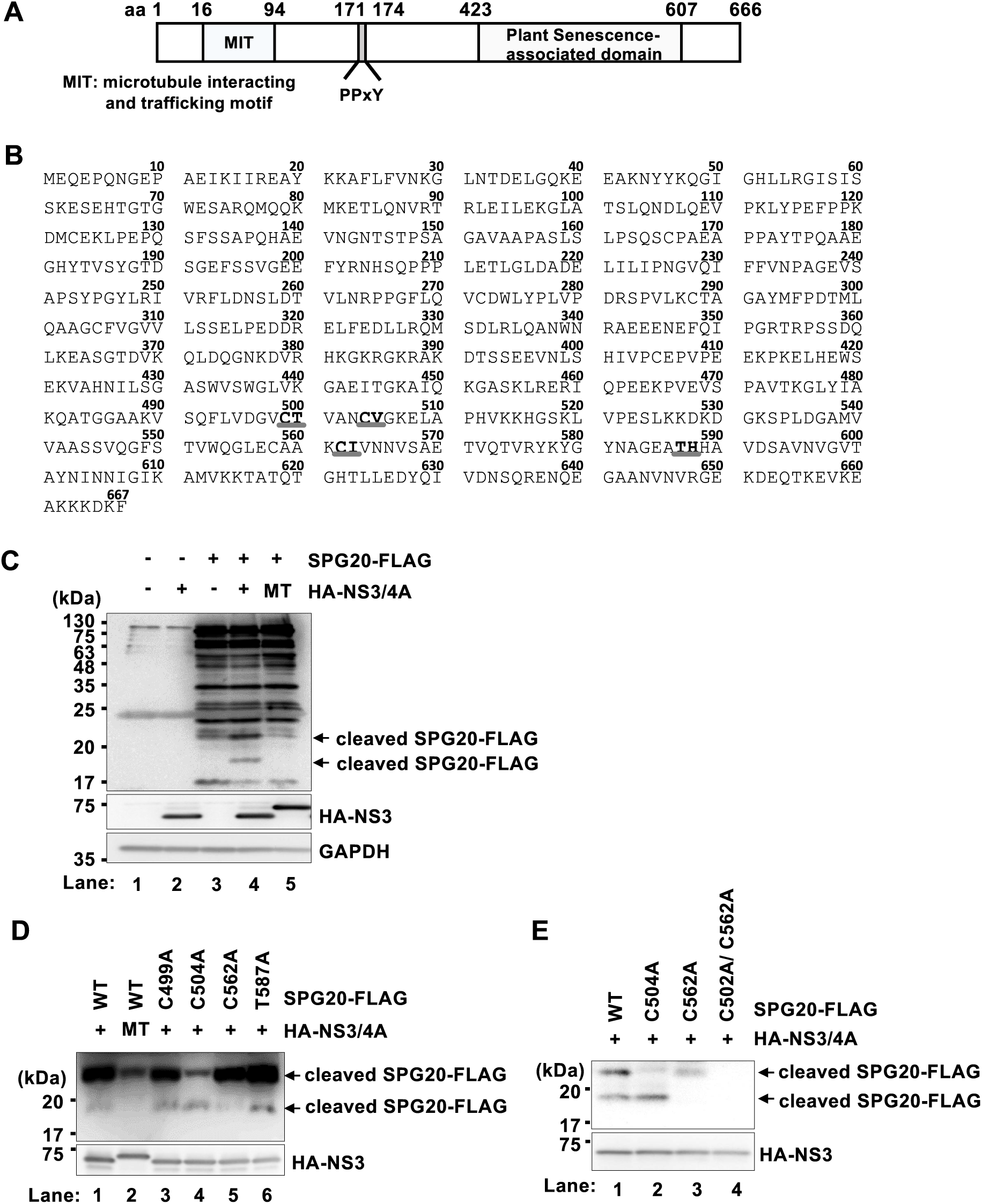
HCV NS3/4A cleaves SPG20 at Cys^504^ and Cys^562^. (A) Schematic representation of SPG20. Microtubule interaction and trafficking motif (MIT) (aa 16 to 94). PPxY motif (aa 171-174). Plant senescence-associated domain (aa 423-607). (B) Amino acid sequence of SPG20. There are at least four possible cleavage sites on SPG20 by HCV NS3/4A (bold, gray underlined). (C) Huh-7.5 cells were plated at 1.5×10^6^ cells per 10 cm dish and cultured for 12 h. The cells were subsequently transfected with pCAG-SPG20-FLAG together with either pCAG-HA-NS3/4A or pCAG-HA-NS3(S139A)/4A. MT, pCAG-HA-NS3(S139A)/4A. At 48 h posttransfection, the cells were harvested. Cell lysates were immunoprecipitated with anti-FLAG M2 beads and bound proteins were immunoblotted with anti-FLAG M2 mouse MAb, anti-HA rabbit PAb, and anti-GAPDH mouse MAb. The level of GAPDH served as a loading control. The arrow indicates cleaved SPG20-FLAG. (D) Huh-7.5 cells were plated at a density of 1.5×10^6^ cells per 10 cm dish and cultured for 12 h. Cells were transfected with each SPG20 mutant plasmid with alanine mutations at Cys^499^, Cys^504^, Cys^562^, or Thr^587^ together with either pCAG-HA-NS3/4A or pCAG-HA-NS3 S139A/4A. MT, pCAG-HA-NS3(S139A)/4A. At 48 h post-transfection, the cells were harvested. The cell lysates were immunoprecipitated with anti-FLAG M2 beads, and the bound proteins were immunoblotted with anti-FLAG MAb and anti-HA PAb. The arrow indicates cleaved SPG20-FLAG. (E) Huh-7.5 cells were transfected with pCAG-SPG20 C504A-FLAG, pCAG-SPG20 C562A-FLAG, or pCAG-SPG20 C504A/C562A-FLAG together with pCAG-HA-NS3/4A. At 48 h post-transfection, the cells were harvested. The cell lysates were immunoprecipitated with anti-FLAG M2 beads, and the bound proteins were immunoblotted with anti-FLAG MAb and anti-HA PAb. The arrow indicates cleaved SPG20-FLAG.

We generated a series of SPG20-FLAG plasmids carrying a substitution mutation at a possible cleavage site. Immunoprecipitation analysis revealed that the 22 kDa fragment of SPG20-FLAG was decreased in the cells expressing SPG20 C504A-FLAG (Fig. 3D, upper panel, lane 4). In addition, the 19 kDa fragment of SPG20-FLAG was markedly decreased in the cells expressing SPG20 C562A-FLAG (Fig. 3D, upper panel, lane 5). On the other hand, two cleaved bands of SPG20 were detected in other mutants, namely SPG20 C499A-FLAG and SPG20 T587A-FLAG (Fig. 3D, upper panel, lanes 3 and 6). These results suggest that the NS3/4A protease cleaves SPG20 at Cys^504^ and Cys^562^. To further examine whether the NS3/4A protease cleaves SPG20 at Cys^504^ and Cys^562^, we generated a plasmid expressing the double mutants SPG20 C504A/C562A-FLAG. Immunoprecipitation analysis revealed that cleaved SPG20-FLAG bands were not detected in the cells expressing C504A/C562A-FLAG (Fig. 3E, upper panel, lane 4). These results indicate that the NS3/4A protease cleaves SPG20 at Cys^504^ and Cys^562^.

### Ubiquitin ligase Itch mediates the polyubiquitylation of ADRP

To determine whether HCV-induced JNK activation results in the phosphorylation of Itch, we examined the phosphorylation of Itch in HCV J6/JFH1-infected cells. The phosphorylation of Itch at Thr222 was increased in HCV-infected cells, indicating that HCV infection induces Itch activation via JNK activation (Fig. 4A, first panel, lanes 2 and 4).

**Fig. 4.**
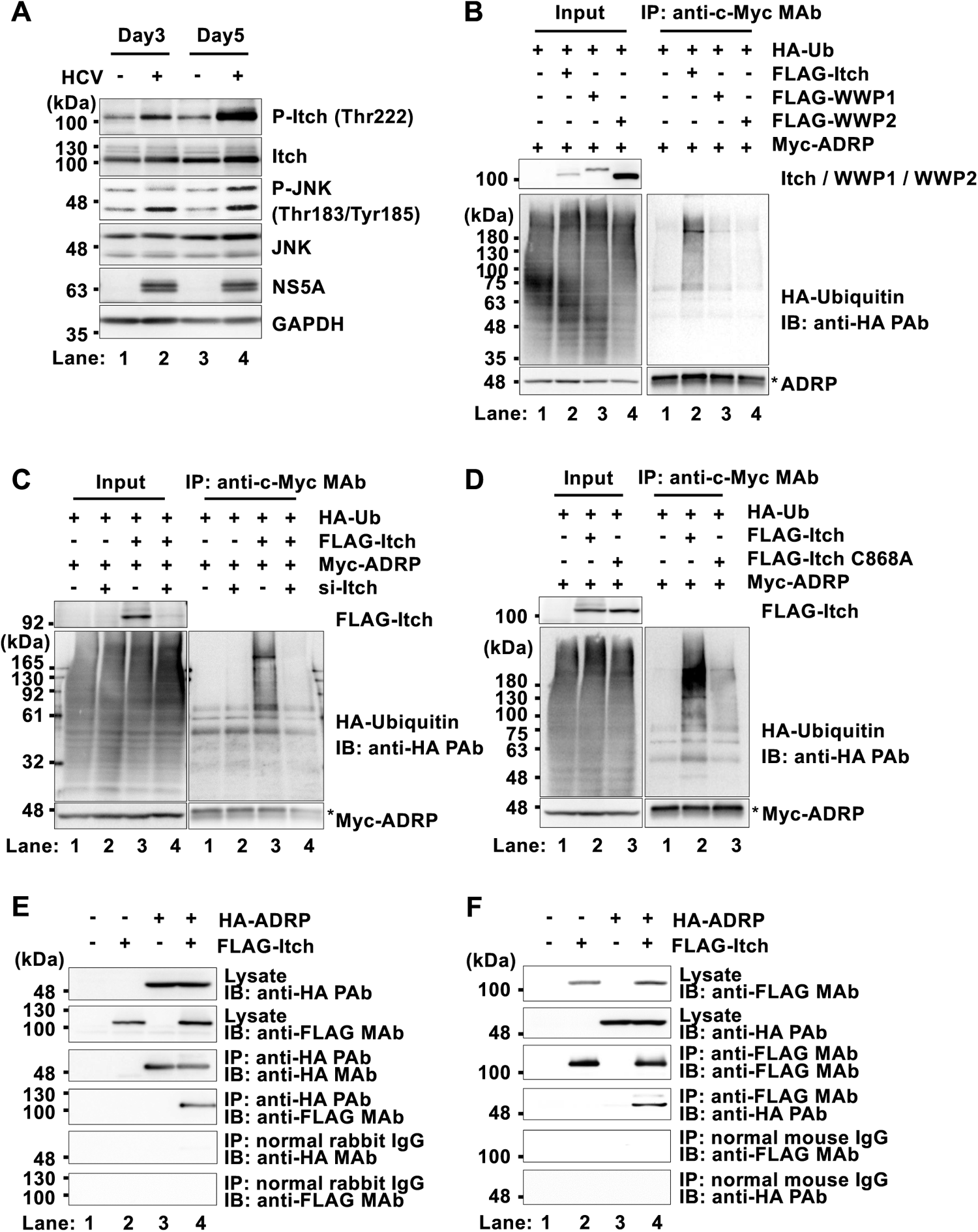
Itch mediates the ubiquitylation of ADRP in Huh-7.5 cells. (A) Huh-7.5 cells were plated in 2.0×10^5^ cells per 6-well plate and cultured for 12 h. The cells were infected with HCV J6/JFH1 at an m.o.i. of 2. The cells were cultured and harvested at the indicated time points and subsequently analyzed by immunoblotting with anti-p-Itch (Thr222) rabbit PAb, anti-Itch rabbit PAb, anti-p-JNK (Thr183/Tyr185) rabbit PAb, anti-JNK rabbit MAb, anti-NS5A rabbit PAb and anti-GAPDH mouse MAb. The level of GAPDH served as a loading control. (B) Huh-7.5 cells were plated at 1.5×10^6^ cells per 10 cm dish and cultured for 12 h. The cells were transfected with pCAG-myc-ADRP and either pCAG-FLAG-Itch, pCAG-FLAG-WWP1, or pCAG-FLAG-WWP2, together with a plasmid encoding HA-tagged ubiquitin. At 48 h after transfection, the cells were harvested. Cell lysates were immunoprecipitated with anti-c-Myc mouse MAb, and bound proteins were immunoblotted with anti-HA rabbit PAb, anti-c-Myc MAb, and anti-FLAG M2 mouse MAb. (C) Huh-7.5 cells were plated at a density of 1.5×10^6^ cells per 10 cm dish and cultured for 12 h. The cells were transfected with either Itch-specific siRNA or universal negative control siRNA. At 24 h after siRNA transfection, cells were transfected with pCAG-myc-ADRP and pCAG-FLAG-Itch, together with a plasmid encoding HA-tagged ubiquitin. At 48 h after transfection, the cells were harvested. Cell lysates were immunoprecipitated with anti-c-Myc MAb, and bound proteins were immunoblotted with anti-HA PAb, anti-c-Myc MAb, and anti-FLAG MAb. (D) Huh-7.5 cells were plated at a density of 1.5×10^6^ cells per 10 cm dish and cultured for 12 h. The cells were transfected with pCAG-Myc-ADRP and either pCAG-FLAG-Itch, or pCAG-FLAG-Itch C868A, together with a plasmid encoding HA-tagged ubiquitin. At 48 h after transfection, the cells were harvested. The cell lysates were immunoprecipitated with anti-c-Myc MAb, and the bound proteins were immunoblotted with anti-HA PAb, anti-c-Myc Mab and anti-FLAG MAb. (E) Huh-7.5 cells were plated at 1.5×10^6^ cells per 10 cm dish and cultured for 12 h. The cells were transfected with pCAG-FLAG-Itch together with pCAG-HA-ADRP. At 48 h after transfection, the cells were harvested. Cell lysates were immunoprecipitated with anti-HA PAb or normal rabbit IgG, and bound proteins were immunoblotted with anti-FLAG MAb and anti-HA mouse MAb. (F) Cell lysates were immunoprecipitated with anti-FLAG MAb or normal mouse IgG, and bound proteins were immunoblotted with anti-FLAG MAb and anti-HA PAb.

To identify an E3 ligase for the ubiuitylation of ADRP in Huh-7.5 cells, Huh-7.5 cells were co-expressed with Myc-ADRP and a plasmid encoding HA-tagged ubiquitin together with either FLAG-Itch, FLAG-WWP1, or FLAG-WWP2. A cell-based ubiquitylation assay revealed that overexpression of FLAG-Itch promoted polyubiquitylation of ADRP protein (Fig. 4B, right, upper panel, lane 2). On the other hand, neither WWP1 nor WWP2 promoted polyubiquitylation of the ADRP protein (Fig. 4B, right, upper panel, lane 3 and 4). In addition, siRNA-mediated knockdown of Itch inhibited the polyubiqutylation of ADRP (Fig. 4C, right, upper panel, lane 4). Furthermore, overexpression of FLAG-Itch increased polyubiquitylation of ADRP, whereas overexpression of the inactive mutant FLAG-Itch C868A failed to promote the polyubiquitylation of ADRP (Fig. 4D, right, upper panel, lanes 2 and 3). These results suggest that Itch specifically mediates the polyubiquituylation of ADRP in Huh-7.5 cells.

To determine whether Itch interacts with ADRP, Huh-7.5 cells were co-expressed with FLAG-Itch together with HA-ADRP. Immunoprecipitation analysis revealed that HA-ADRP was coimmunoprecipitated with FLAG-Itch using anti-HA PAb (Fig. 4E, fourth panel, lane 4). Conversely, immunoprecipitation analysis revealed that HA-ADRP was coimmunoprecipitated with FLAG-Itch using anti-FLAG MAb (Fig. 4F, fourth panel, lane 4). These results suggest that Itch interacts with ADRP in Huh-7.5 cells. We concluded that the ubiquitin ligase Itch mediates the polyubiquitylation of the ADRP protein in Huh-7.5 cells.

### HCV infection induces the ubiquitin-dependent proteasomal degradation of ADRP

To determine whether HCV infection affects the expression levels of the ADRP protein, we performed immunoblot analysis of the endogenous ADRP protein in HCV J6/JFH1-infected Huh-7.5 cells. Immunoblot analysis revealed that the protein level of ADRP was markedly lower in HCV-infected cells than in mock-infected control cells (Fig. 5A, first panel, lanes 2, 4 and 6). On the other hand, TIP47 protein levels did not change (Fig. 5A, second panel, lanes 2, 4 and 6). The proteasome inhibitor MG132 restored the protein level of ADRP (Fig. 5B, first panel, lanes 6 and 8). These results suggest that HCV-infection promotes proteasomal degradation of the ADRP protein.

**Fig. 5.**
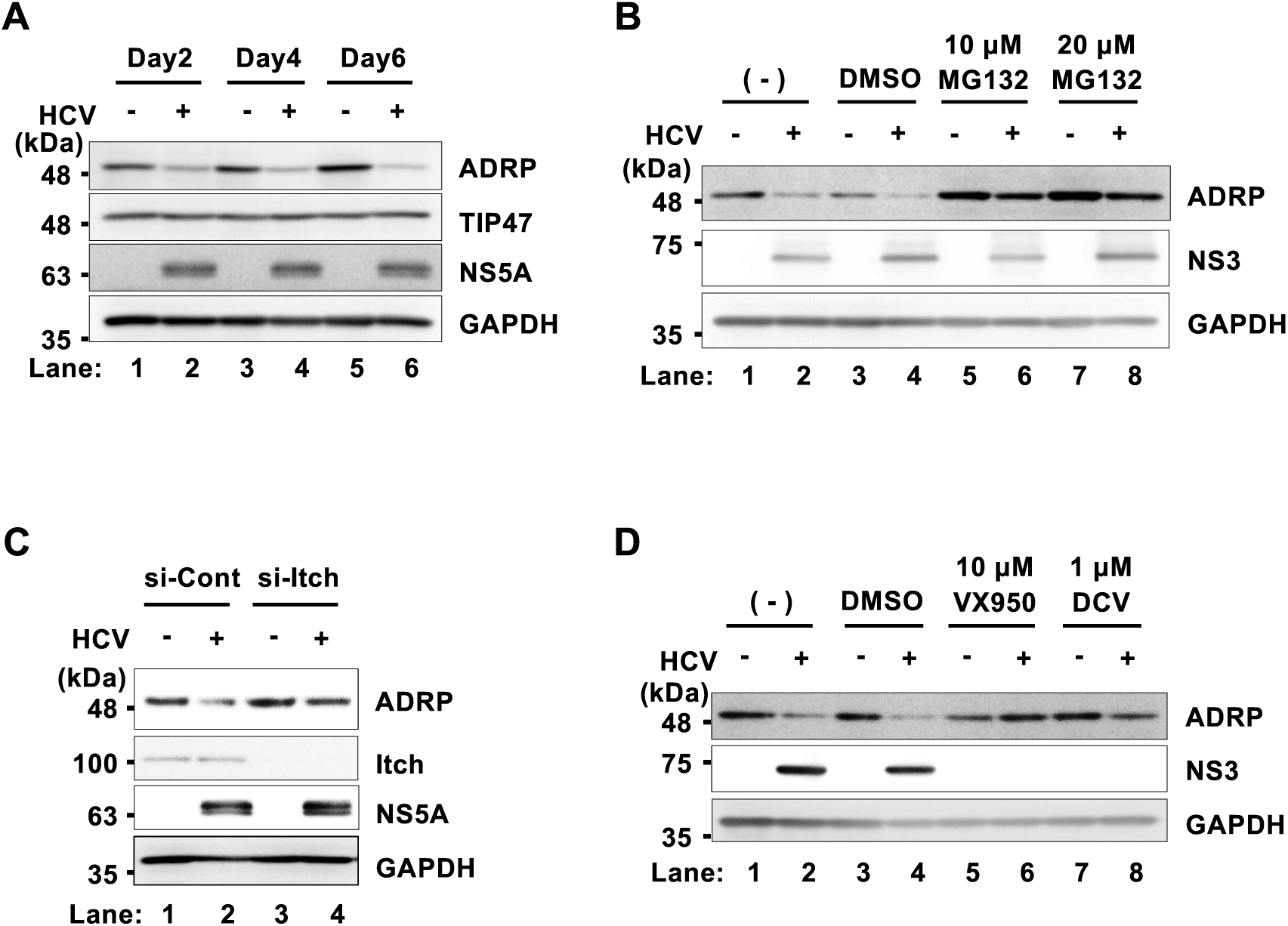
HCV infection promotes Itch-mediated degradation of ADRP via the ubiquitin‒proteasome pathway. (A) Huh-7.5 cells were plated in 2.0×10^5^ cells per 6-well plate and cultured for 12 h. The cells were infected with HCV J6/JFH1 at an m.o.i. of 2. The cells were then cultured and harvested at the indicated time points. The cells were analyzed by immunoblotting with anti-ADRP mouse MAb, anti-TIP47 mouse MAb, anti-NS5A rabbit PAb and anti-GAPDH mouse MAb. The level of GAPDH served as a loading control. (B) Huh-7.5 cells were plated in 2.0×10^5^ cells per 6-well plate and cultured for 12 h. The cells were infected with HCV J6/JFH1 at an m.o.i. of 2. At 5 days post-infection, 10 μM or 20 μM MG132 proteasome inhibitor was administered to the cells. The cells were cultured for 12 h and harvested. The cell lysates were analyzed by immunoblotting with anti-ADRP MAb, anti-NS3 MAb and anti-GAPDH MAb. (C) Huh-7.5 cells were plated in 3.0×10^5^ cells per 12-well plate. The cells were transfected with either Itch-specific siRNA or universal negative control siRNA and cultured. At 24 h after siRNA transfection, the cells were infected with HCV J6/JFH1 at an m.o.i. of 2, cells were cultured for 48 h and harvested. The cell lysates were analyzed by immunoblotting with anti-ADRP MAb, anti-Itch mouse MAb, anti-NS5A PAb and anti-GAPDH MAb. (D) Huh-7.5 cells were plated in 2.0×10^5^ cells per 6-well plate and cultured for 12 h. The cells were infected with HCV J6/JFH1 at an m.o.i. of 2. At 4 h postinfection, 10μM VX950 or 1 μM daclatasvir was administered to the cells. The cells were cultured for 5 days after infection and harvested. The cell lysates were analyzed by immunoblotting with anti-ADRP MAb, anti-NS3 MAb, and anti-GAPDH MAb.

To determine whether Itch plays a role in HCV-induced proteasomal degradation of the ADRP protein, Itch was knocked down by siRNA (Fig. 5C). siRNA-mediated knockdown of Itch restored the level of ADRP in HCV-infected cells (Fig. 5C, first panel, lane 4). These results suggest that HCV infection promotes Itch-mediated ubiquitin-dependent degradation of ADRP.

To determine whether HCV is involved in the degradation of ADRP, we assessed the effects of the NS3/4A protease inhibitor VX950 or the NS5A inhibitor DCV on the levels of ADRP in HCV-infected cells. Treatment of HCV-infected cells with either VX950 or daclatasvir restored the protein levels of ADRP (Fig. 5D, first panel, lanes 6 and 8). These results suggest that HCV infection induces the degradation of ADRP.

### ADRP accumulates around LDs in HCV-infected cells

To examine the subcellular localization of ADRP in HCV-infected Huh-7.5 cells, we performed immunofluorescence staining with a BODIPY lipid probe. In mock-infected cells, ADRP was localized in both the nucleus and cytoplasm (Fig. 6, upper panel, ADRP). However, ADRP was localized mainly on the LD surface in HCV-infected cells (Fig. 6, lower panel, Merge). These results suggest that HCV infection promotes the accumulation of ADRP on the surface of LDs.

**Fig. 6.**
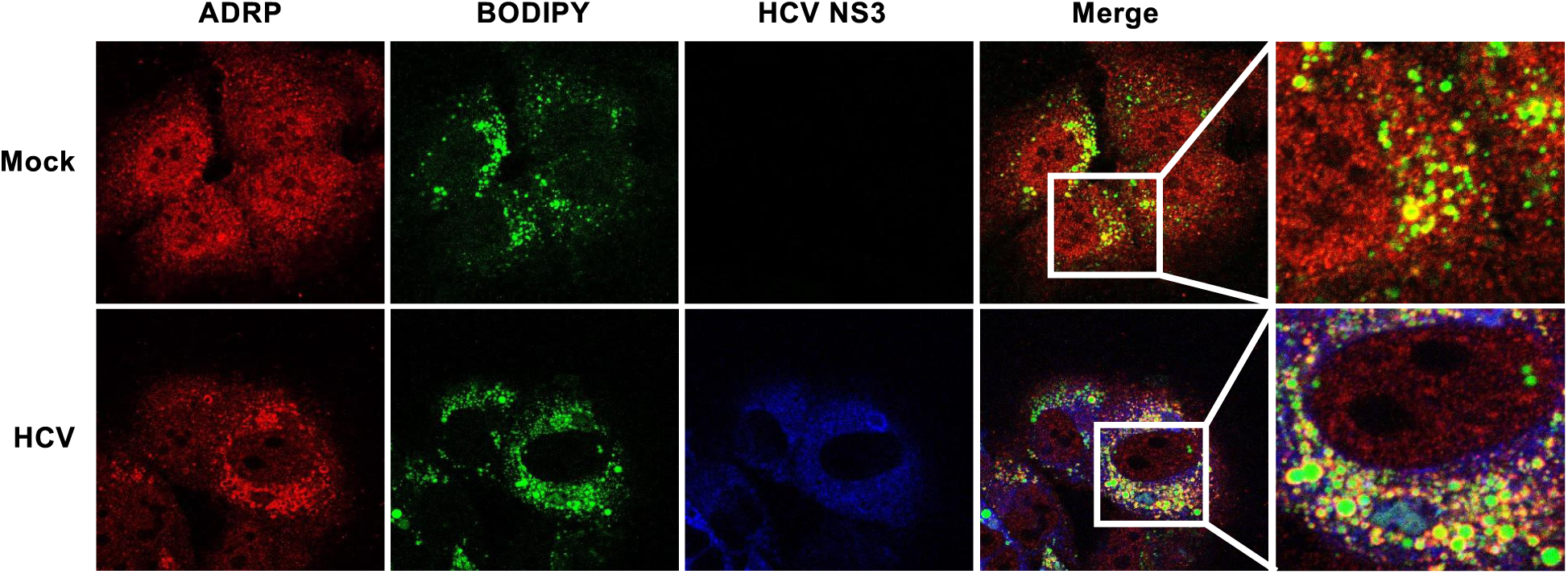
ADRP accumulates around lipid droplets in HCV-infected cells. Huh-7.5 cells were plated in 1.0×10^5^ cells per 24-well plate and cultured for 12 h. The cells were infected with HCV J6/JFH1 at an m.o.i. of 2. At 6 days post-infection, the cells were fixed with 4% paraformaldehyde and stained with anti-ADRP rabbit PAb followed by Alexa Fluor 594-conjugated goat anti-rabbit IgG (red), and anti-NS3 MAb followed by Alexa Fluor 405-conjugated goat anti-mouse IgG (blue). Lipid droplets were stained with BODIPY493/503 (green). The stained cells were examined under a confocal laser scanning microscope.

### NS3/4A protease cleaves SPG20 and inhibits the SPG20‒TIP47 interaction

We investigated why ADRP proteins on the surface of LDs are not degraded by the ubiquitin-proteasome pathway in HCV-infected cells, even though the Itch-mediated ubiquitin-dependent degradation of ADRP is promoted. SPG20 is known to interact with TIP47 at the surface of LDs. The region raging from aa 433 to aa 584 on SPG20 (Fig. 3A) was found to be important for the interaction with TIP47 (26). Therefore, we hypothesize that the NS3/4A protease cleaves SPG20 at Cys^504^ and Cys^562^, resulting in the inhibiton of the SPG20**‒**TIP47 interaction, thereby disrupting the recruitment of Itch to LD-attached ADRP (Fig.7A).

**Fig. 7.**
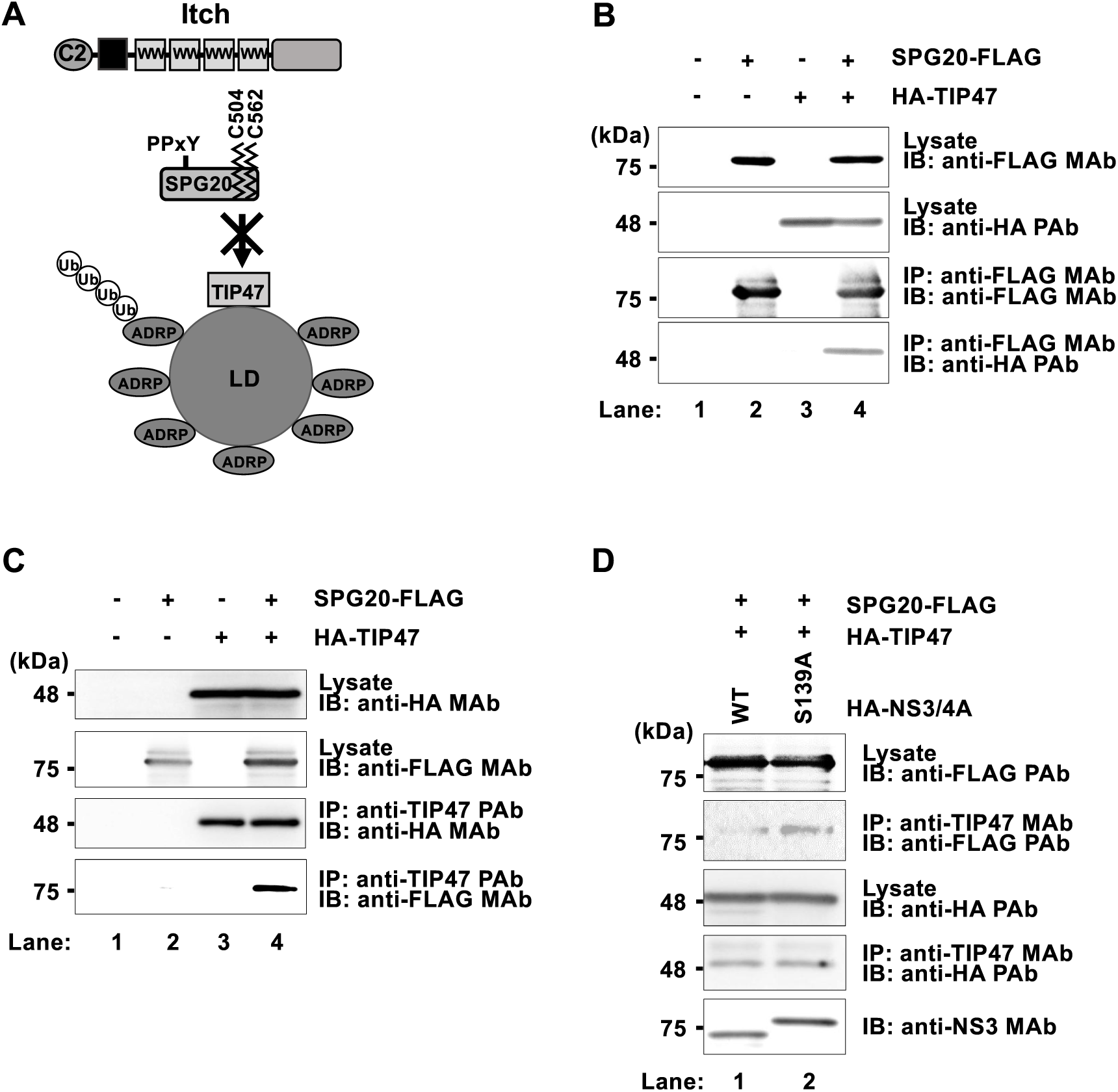
HCV NS3/4A protease cleaves SPG20 and inhibits the SPG20-TIP47 interaction. (A) Working hypothesis: the HCV NS3/4A protease cleaves SPG20 at Cys^504^ and Cys^562^. When SPG20 is cleaved by NS3/4A, SPG20 cannot interact with TIP47, and Itch is not recruited to LDs. (B) Huh-7.5 cells were plated in 1.5×10^6^ cells per 10 cm dish and cultured for 12 h. The cells were then transfected with pCAG-SPG20-FLAG together with pCAG-HA-TIP47. At 48 h after transfection, the cells were harvested. The cell lysates were immunoprecipitated with anti-FLAG M2 beads, and the bound proteins were immunoblotted with anti-FLAG M2 mouse MAb and anti-HA rabbit PAb, respectively. (C) Cell lysates were immunoprecipitated with anti-TIP47 PAb, and bound proteins were immunoblotted with anti-FLAG MAb and anti-HA MAb, respectively. (D) Huh-7.5 cells were transfected with pCAG-HA-NS3/4A or pCAG-HA-NS3(S139A)/4A together with pCAG-SPG20-FLAG and pCAG-HA-TIP47. The cell lysates were immunoprecipitated with anti-TIP47 MAb, and the bound proteins were immunoblotted with anti-FLAG MAb, anti-HA PAb, or anti-NS3 MAb.

To determine whether SPG20 interacts with TIP47 in Huh-7.5 cells, we transfected cells with pCAG-SPG20-FLAG together with pCAG-HA-TIP47. Immunoprecipitation analysis revealed that HA-TIP47 was coimmunoprecipitated with SPG20-FLAG using anti-FLAG MAb (Fig. 7B, right panel, fourth panel, lane 4). Conversely, immunoprecipitation analysis revealed that SPG20-FLAG was coimmunoprecipitated with HA-TIP47 using anti-TIP47 PAb (Fig. 7C, fourth panel, lane 4). These results suggest that SPG20 interacts with TIP47 in Huh-7.5 cells.

To further confirm that the NS3/4A protease cleaves SPG20 and inhibits SPG20**‒** TIP47 interaction, we performed immunoprecipitation analysis. Immunoprecipitation analysis revealed that the SPG20-FLAG did not co-immunoprecipitate with HA-TIP47 in the presence of wild-type NS3/4A (Fig. 7D, second panel, lane 1). However, SPG20-FLAG was co-immunoprecipitated with HA-TIP47 in the presence of inactive NS3/4A mutant (Fig. 7D, second panel, lane 2). These results suggest that the NS3/4A protease cleaves SPG20, thereby inhibiting the SPG20**‒**TIP47 interaction.

Taken these results together, we propose a model in which the HCV NS3/4A protease cleaves SPG20 at Cys^504^ and Cys^562^, and inhibits the SPG20**‒**TIP47 interaction, thereby inhibiting the recruitment of Itch to LD-attached ADRP, preventing the ubiquitin-dependent degradation of LD-attached ADRP and promoting the formation of large LDs (Fig. 8).

**Fig. 8.**
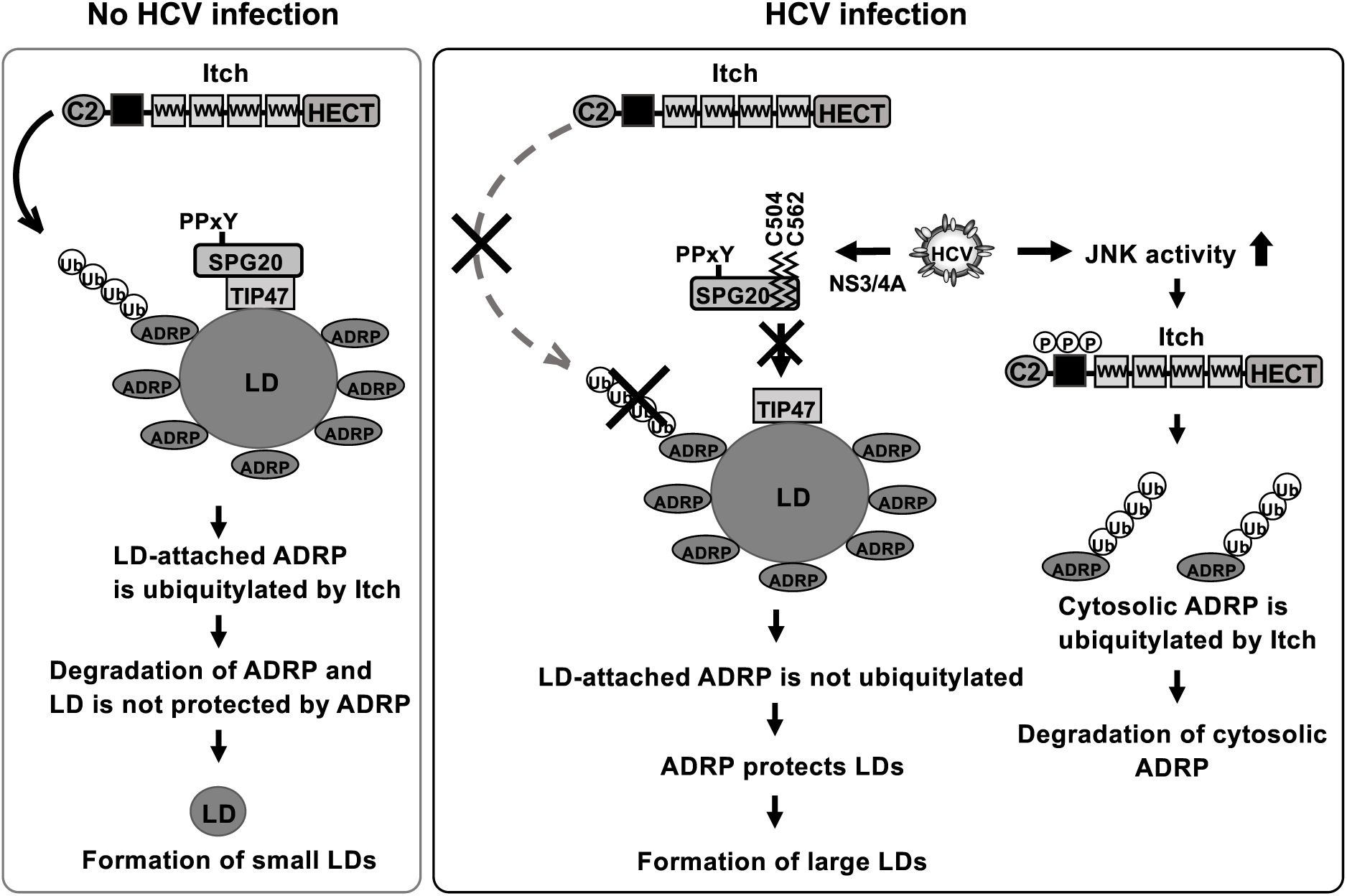
A proposed model of HCV-induced large LDs formation via the cleavage of SPG20 by HCV NS3/4A protease. Left, no HCV infection: ADRP localizes on the surface of LDs and protects LDs from lipase-mediated degradation. SPG20 localizes to the surface of LDs via the association of TIP47. Itch ubiquitin ligases are recruited to the surface of LDs via interactions between the WW domain of Itch and the PPxY motif of SPG20. Itch mediates the polyubiquitylation of ADRP and promotes proteasomal degradation. LDs lack ADRP on their surface and decrease in size. Right, HCV infection: When cells are infected with HCV, HCV infection increases JNK activity, resulting in the phosphorylation and activation of Itch ubiquitin ligases. Active Itch promotes polyubiquitylation of cytosolic ADRP and promotes proteasomal degradation. The HCV NS3/4A protease cleaves SPG20 at Cys^504^ and Cys^562^. When SPG20 is cleaved by NS3/4A, SPG20 cannot interact with TIP47, and Itch is not recruited to LDs. LD-attached ADRP is not ubiquitylated and lacks ubiquitin-dependent degradation. Therefore, ADRP remains on the surface of LDs and protects them, leading to the formation of large LDs in HCV-infected cells.

## DISCUSSION

LDs play important roles in HCV RNA replication and the production of HCV particles (6, 27). HCV infection often causes the accumulation of large LDs in hepatocytes. In this study, we aimed to elucidate the molecular mechanism underlying large LD formation induced by HCV infection. Here, we demonstrated that HCV infection promoted the formation of large LDs (Fig. 1A-1D). However, the number of LDs per cell was not increased in HCV-infected cells (Fig. 1E). We also demonstrated that SPG20 was cleaved in HCV-infected cells (Fig. 2A). SPG20 was cleaved in the presence of the HCV NS3/4A protease (Fig. 2C) but not the HCV NS2 protease (Fig. 2B). An inactive mutant of the HCV NS3/4A protease abolished the cleavage of SPG20 (Fig. 2D). These findings suggest that the HCV NS3/4A protease is responsible for the cleavage of SPG20. On the basis of the consensus sequence for the HCV NS3/4A protease cleavage site, there were at least four possible cleavage sites on SPG20 by the HCV NS3/4A protease (Fig. 3B). Site-directed mutagenesis revealed that Cys^504^ and Cys^562^ of SPG20 are important for cleavage by the HCV NS3/4 protease (Fig. 3D and 3E). Cys^504^ and Cys^562^ of SPG20 reside within the SPG20**‒**TIP47 interaction region. Therefore, we hypothesized that HCV NS3/4A cleaves SPG20 at Cys^504^ and Cys^562^, thereby inhibiting the interaction between SPG20 and TIP47. The NS3 protein has been detected at the LD surface (8, 28), consistent with the notion that the HCV NS3/4A protease cleaves the LD-associated protein SPG20.

ADRP plays an important role in LD biogenesis. SPG20 is known to play a key role in the turnover of lipid droplets (14, 16). A recent study (29) demonstrated that SPG20 functions as a lipophagy receptor. SPG20 inhibition in cultured human neurons or murine brain neurons enhanced LD formation, suggesting that impaired LD metabolism is involved in the development of Troyer syndrome.

Nedd4 family E3 ligases contain WW domain (30). This domain is responsible for substrate recognition via the PPxY motif. Hopper C et al. (16) reported that SPG20 interacts with the WW domains of Itch via the PPxY motif of SPG20. The interaction between SPG20 and Itch increases the enzymatic activity of the E3 ligase of Itch. SPG20 is an adopter of Itch for the polyubiquitylation of the LD-attached ADRP. We demonstrated that Itch promoted polyubiquitylation of ADRP in Huh-7.5 cells. However, neither WWP1 nor WWP2 promoted the polyubiquitylation of ADRP (Fig. 4B). ADRP was co-immunoprecipitated with Itch (Fig. 4E and 4F). These data suggest that Itch is responsible for the ubiquitin-dependent degradation of ADRP in Huh-7.5 cells.

Itch is directly activated by phosphorylation of the proline-rich region at S199, S232, and T222 via JNK protein kinase and conformational changes (31). Itch phosphorylation is important for the interaction between the WW domain and HECT domain of Itch, which leads to a conformational change from the closed inactive form to the open active form. This conformational change in Itch is important for the polyubiquitylation of its substrates. We previously reported that Itch promotes the release of HCV particles via polyubiquitylation of VPS4A, leading to the activate of the ROS/JNK/Itch/VPS4A signaling pathway for the release of infectious HCV particles (25). We confirmed that Itch activation with its phosphorylation at Thr222 was induced via JNK activation in HCV-infected cells (Fig. 4A).

The protein levels of ADRP were decreased in HCV J6/JFH1-infected cells (Fig. 5A). Treatment of the cells with MG132, a proteasome inhibitor, restored the protein levels of ADRP (Fig. 5B), suggesting that the protein level of ADRP is decreased in HCV-infected cells via the ubiquitin-proteasome pathway. In mock-infected cells, ADRP was localized in both the cytoplasm and the nucleus (Fig. 6). On the other hand, ADRP was co-localized with the surface of LDs in HCV-infected cells. We provided evidence that NS3/4A-mediated cleavage of SPG20 inhibited SPG20**‒**TIP47 interaction, thereby disrupting the recruitment of Itch to LDs (Fig. 7). We propose that LD-attached ADRP can prevent ubiquitin-dependent degradation and protect LDs, thereby promoting the formation of larger LDs (Fig. 8).

LD-binding proteins are classified into two categories, as summarized by Kory et al. (32). Class I proteins, such as oleosin and glycerol-3-phosphate acyltransferase 4 (GPAT4), are translated at the ER and localize to both the ER and the LD surface. Class II proteins, such as PAT (perilipin/ADRP/TIP47)-domain proteins, harbor a central 11-mer repeat-containing domain that is predicted to fold into amphipathic and hydrohobic helices upon membrane binding. The C-terminal four-helix bundle of ADRP and TIP47 participates in LD binding. In this study, we clarified the role of ADRP in the formation of large LDs.

In conclusion, we propose a novel mechanism in which the HCV NS3/4A protease cleaves SPG20 and inhibits the SPG20**‒**TIP47 interaction, thereby preventing the ubiquitin-dependent degradation of ARDP on the surface of LDs and promoting the formation of large LDs in HCV-infected cells. Understanding the molecular mechanism underlying the HCV-induced formation of large LDs may shed new light on the treatment of steatosis and chronic liver diseases caused by HCV infection.

## MATERIALS AND METHODS

### Cell culture

The human hepatoma cell line Huh-7.5 (33) was kindly provided by Dr. Charles M. Rice (The Rockefeller University, NY). The cells were cultured in Dulbecco’s modified Eagle’s medium (DMEM) (High Glucose) with L-glutamine (Wako, Osaka, Japan) supplemented with 50 IU/ml penicillin, 50 µg/ml streptomycin (Gibco, Grand Island, NY), 10% heat-inactivated fetal bovine serum (Biowest, Nuaillé, France), and 0.1 mM nonessential amino acids (Invitrogen, New York, NY) at 37°C in a 5% CO_2_ incubator. The cells were transfected with plasmid DNA via FuGene6 transfection reagents (Promega, Madison, WI).

The pFL-J6/JFH1 plasmid, encoding the entire viral genome of a chimeric strain of HCV-2a, JFH1 (34), was kindly provided by Charles M. Rice. The HCV genomic RNA was synthesized *in vitro* via the use of pFL-J6/JFH1 as a template and was transfected into Huh-7.5 cells by electroporation (35–37). The virus produced in the culture supernatant was used for infection experiments (35).

### Expression plasmids

To express NS3/4A, the cDNA fragment of nt 3420 to 5474 derived from the HCV Con1 strain was amplified by polymerase chain reaction (PCR). The specific primers used for PCR were as follows: sense primer, 5’-TAACTCGAGCGCGCCTATTACGGCCTACTC-3’; antisense primer, 5’-AAAGCGGCCGCTCAGCACTCTTCCATCTCATCGA-3’. The amplified PCR product was purified, cloned, and inserted into the XbaI-NotI site of pCAG-HA. The plasmids pCAG-FLAG-NS2-3ΔC and pCAG-HA-NS3(S139A)/4A were kindly provided by Dr. Suzuki (Hamamatsu University School of Medicine, Shizuoka, Japan). The plasmids pCAG-FLAG-SPG20, pCAG-FLAG-Itch, pCAG-FLAG-WWP2, pCAG-HA-ADRP, and pCAG-HA-TIP47 were constructed by amplification of cDNA fragments by PCR. The cDNA fragments were amplified via the use of pBluescriptR-SPG20, pCMV-SPORT6-WWP2, pOTB7-ADRP, and pDNR-Dual-TIP47 as templates. These plasmids were purchased from the PlasmID database (Harvard Medical School, MA). The expression plasmid pCAG-FLAG-WWP1 was previously described (38). The specific primers used for PCR were as follows: SPG20 sense primer, 5’-TCGAGCTCAGCGGCCGCCATGGAGCAAGAGCCACAA-3’; SPG20 antisense primer, 5’-AGTGAATTCGCGGCCGCTCATTTATCTTTCTTCTT-3’, WWP1 sense primer, 5’-TCGAGCTCAGCGGCCGCCATGGCCACTGCTTCACCA-3’; WWP1 antisense primer, 5’-AGTGAATTCGCGGCCGCTCATTCTTGTCCAAATCC-3’, WWP2 sense primer, 5’-TCGAGCTCAGCGGCCATGGCATCTGCCAGCTCT-3’; WWP2 antisense primer, 5’-AGTGAATTCGCGGCCTTACTCCTGTCCAAAGCCC-3’, ADRP sense primer, 5’-TCGAGCTCAGCGGCCGCCATGGCATCCGTTGCAGTT-3’; ADRP antisense primer, 5’-AGTGAATTCGCGGCCGCTTAATGAGTTTTATGCTC-3’, TIP47 sense primer, 5’-TCGAGCTCAGCGGCCGCCATGTCTGCCGACGGGGC-3’; TIP47 antisense primer, 5’-AGTGAATTCGCGGCCGCCTACTTCTTCTCCTCCGG-3’.

These amplified PCR products were subsequently purified. Each of them was inserted into the Not I site of pCAG-FLAG or pCAG-HA via an In-Fusion HD-Cloning Kit (Clontech, Mountain View, CA). To express ADRP as a myc-tagged protein, the cDNA fragment was amplified by PCR, cloned, and inserted into the Kpn I/Bgl II site of pCAG-MCS2. The specific primers used for PCR were as follows: ADRP sense primer, 5’-AAGGTACCATGGAGCAAAAGCTCATTTCTGAAGAGGACTTGATGGCATCCGTTGCAGTTGAT-3’; ADRP antisense primer, 5’-TTAGATCTTTAATGAGTTTTATGCTCAGATCG-3’.

The cDNA fragment for the pCAG-SPG20-FLAG plasmid was amplified by PCR, cloned, and inserted into the Not I site of pCAG-MCS2. The specific primers used for PCR were as follows: SPG20-FLAG sense primer, 5’-AAGCGGCCGCACCATGGAGCAAGAGCCACAAAAT-3’; SPG20-FLAG antisense primer, 5’-TTGCGGCCGCTTACTTATCGTCGTCATCCTTGTAATCTTTATCTTTCTTCTTTGC CTC-3’.

The point mutations of SPG20 were introduced into pCAG-SPG20-FLAG via overlap extension using PCR. The specific primers used for PCR were as follows: SPG20 C499A sense primer, 5’-CTGGTTGATGGAGTTGCCACTGTAGCAAATTGC-3’; SPG20 C499A antisense primer, 5’-GCAATTTGCTACAGTGGCAACTCCATCAACCAG-3’ (C499A). SPG20 C504A sense primer, 5’-TGCACTGTAGCAAATGCCGTTGGAAAAGAA-3’; SPG20 C504A antisense primer, 5’-TTCTTTTCCAACGGCATTTGCTACAGTACG-3’. SPG20 C562A sense primer, 5’-GAATGTGCAGCTAAAGCCATCGTTAACAAT-3’; SPG20 C562A antisense primer, 5’-ATTGTTAACGATGGCTTTAGCTGCACATTC-3’, SPG20 T587A sense primer, 5’-AATGCAGGAGAAGCTGCCCACCATGCGGTG-3’; SPG20 T587A antisense primer, 5’-CACCGCATGGTGGGCAGCTTCTCCTGCATT-3’. The sequences of the inserts were extensively confirmed by sequencing (Eurofins Genomics, Tokyo, Japan).

### Antibodies

The mouse monoclonal antibodies (MAbs) used in this study were anti-FLAG (M2) MAb (F3165, Sigma, St. Louis, MO), anti-HA MAb (H-3663, Sigma), anti-NS3 MAb (MAB8691, Millipore, Billerica, MA), anti-c-Myc MAb (9E10, Santa Cruz Biotechnology, Santa Cruz, CA), anti-TIP47 MAb (B-3, Santa Cruz Biotechnology), anti-ADRP MAb (03-610102, ARP American Research Products, Waltham, MA), anti-Itch mouse MAb (611198, BD Biosciences, Franklin Lakes, NJ) and anti-glyceraldehyde-3-phosphate dehydrogenase (GAPDH) MAb (MAB374, Millipore). The rabbit MAbs used in this study was anti-SAPK/JNK (56G8) rabbit MAb (9258, Cell Signaling Technology, Beverly, MA). The rabbit polyclonal antibody (PAbs) used in this study were anti-HA PAb (H-6908, Sigma), anti-DDDDK tag (FLAG) PAb, anti-ADFP (ADRP) PAb (ab108323, Abcam, Cambridge, MA), anti-Perilipin3 (TIP47) PAb (ab47639, abcam), anti-Itch rabbit PAb (SAB4200036, Sigma), anti-phospho-Itch (Thr222) rabbit PAb (AB10050, MIllipore) and anti-phospho-SAPK/JNK (Thr183/Tyr185) rabbit PAb (9251, Cell Signaling Technology). The goat PAb used in this study was anti-SPG20 goat PAb (sc-49521, Santa Cruz Biotechnology). Horseradish peroxidase (HRP)-conjugated antibodies, including anti-mouse IgG and anti-rabbit IgG (both from Cell Signaling Technology) as well as donkey anti-goat IgG (Santa Cruz Biotechnology) were used.

### Immunoblot analysis

Immunoblot analysis was performed essentially as described previously (36, 38, 39). The cell lysates were separated by 10% or 15% sodium dodecyl sulfate**‒**polyacrylamide gel electrophoresis (SDS-PAGE) and transferred to polyvinylidene difluoride membrane (Millipore). The membranes were incubated with primary antibodies, followed by incubation with peroxidase-conjugated secondary antibody. The positive bands were visualized using enhanced chemiluminescence (ECL) Western blotting detection reagents (GE Healthcare, Buckinghamshire, UK).

### Immunoprecipitation

Cultured cells were lysed with a buffer containing 150 mM NaCl, 10 mM Tris-HCl (pH 7.4), 0.1% SDS, 1% sodium deoxycholic acid, 1% Triton X-100, and cOmplete^TM^ Protease Inhibitor Cocktail Tablets (Roche Diagnostics, Indianapolis, IN). The lysate was centrifuged at 12,500 × g for 15 min at 4°C and the supernatant was immunoprecipitated with appropriate antibodies. Immunoprecipitation was performed as described previously (40). Briefly, the cell lysates were immunoprecipitated with anti-FLAG M2 affinity gel (Sigma) or protein A-Sepharose 4 fast flow (GE Healthcare) and incubated with appropriate antibodies at 4°C overnight. After being washed with lysis buffer five times, the immunoprecipitates were analyzed by immunoblotting.

### siRNA transfection

HCV-infected Huh-7.5 cells or uninfected control cells (3.0×10^5^ cells per12-well plate) were transfected with 20 pmol of either Itch-specific siRNA duplexes (Qiagen, Valencia, CA) or MISSION siRNA Universal Negative Control (SIC-001, Sigma Genosys) using Lipofectamine RNAiMAX transfection reagent (Life Technologies, Carlsbad, CA) according to the manufacture’s instructions and cultured for 48 h. The Itch-siRNA target sequences was as follows: 5’-ATGGGTAGCCTCACCATGAUU-3’.

### Immunofluorescence staining

Huh-7.5 cells cultured on glass cover slips were fixed with 4% paraformaldehyde at room temperature for 15 min. After being washed with PBS, the cells were permeabilized with PBS containing 0.1% Triton X-100 for 15 min at room temperature and incubated in PBS containing 1% bovine serum albumin for 60 min to block nonspecific reactions. The cells were incubated with Can Get Signal Immunostain Solution A (TOYOBO, Osaka, Japan) containing mouse anti-NS3 antibody (Millipore) at room temperature for 60 min. The cells were washed four times with PBS and incubated with Can Get Signal Immunostain Solution A containing Alexa Fluor 405-conjugated anti-mouse immunoglobulin G (IgG) (Life Technologies) with BODIPY 493/503 (Molecular Probes, Eugene, OR) at room temperature for 60 min. The cells were washed four times with PBS and then observed under an LSM700 confocal laser scanning microscope (Carl Zeiss, Oberkochen, Germany).

### Quantification and measurement of LD

To analyze the size and the number of LDs, the cells were analyzed with an LSM700 confocal microscope. The cells selected as region of interest (ROIs) were processed by thresholding. ROIs were generated via a free-hand selection tool. The images were used to quantify the LD size per cell. The LDs per cell were counted from the same images. The size and the number of LDs were analyzed via ImageJ (Ver. 1.48).

### Statistics

The results are expressed as the mean ± standard error. Statistical significance was determined by Student’s *t*-test. P-values <0.01 (**) were considered significant.

## Data Availability Statement

All data are presented in the main figures. The data that support the findings of this study is available at bioRxiv (https://www.biorxiv.org/). Raw sequencing data, microscopy images, materials, and sequence information are available upon request. Correspondence and requests for materials should be addressed to Professor Ikuo Shoji.

## ACKNOWLEDGMENTS

We are grateful to C. M. Rice (Rockefeller University, New York, NY) for providing us Huh-7.5 cells and pFL-J6/JFH1. We thank Y. Kozaki for secretarial work. This research was supported by Basic and Clinical Research on Hepatitis from Japan Agency for Medical Research and Development, AMED, under grant number JP18fk021006, JP20fk0210040, JP20fk0210053, and JP21fk0210090. This work was also supported by research grants of the Japan Society for the Promotion of Science (KAKENHI), under grant numbers 19K16671, 20K07514, and 22K15470.

## AUTHOR CONTRIBUTIONS

C.M., and I.S. conceived and designed the experiments. C.M. carried out most of the experiments. T.A. P.Y. and L.D. assisted the constructions and the data analysis. C.M., and I.S. wrote the manuscript. All the authors contributed to the manuscript and approved the submitted version.

